# Scalable data harmonization for single-cell image-based profiling with CytoTable

**DOI:** 10.1101/2025.06.19.660613

**Authors:** Dave Bunten, Jenna Tomkinson, Erik Serrano, Michael J. Lippincott, Kenneth I. Brewer, Vince Rubinetti, Faisal Alquaddoomi, Gregory P. Way

## Abstract

High-content imaging (HCI) involves the automated acquisition and quantitative analysis of cell phenotypes from microscopy images. These studies often rely on screening, which can involve thousands of chemical or genetic perturbations that produce terabytes of microscopy data. To extract meaningful biological insights, this data must be processed into quantitative features through a technique known as image-based profiling. A major analytical bottleneck is curating the high-dimensional, single-cell data derived from varied image analysis tools. These datasets suffer from inconsistent schemas, inefficient file formats, and undocumented ontological relationships. These challenges reduce reproducibility and slow progress in downstream applications. To solve these issues, we introduce CytoTable^1^, a software package for harmonizing single-cell image-based profiling. CytoTable enables modular, portable, and cross-language data integration through a robust, reproducible, and scalable engine that harmonizes single-cell readouts from multiple image analysis tools, preparing for feature integration with software in the Cytomining ecosystem such as Pycytominer.^2^

## Introduction

Image-based profiling is a critical part of bioinformatics processing for high-content imaging (HCI) experiments. HCI uses a pipeline of software tools: each tool reads and processes the output of the one before it, then passes its results on to the next stage (**Fig. 1A**).^3^ These data processing pipelines start with wet lab assays and microscopy imaging devices that produce microscopy images, which are then analyzed with image analysis tools to produce structured, high-content morphology feature measurements. Scientists then use the feature data to find patterns and test hypotheses on specific biological processes and the impact of perturbations on cell phenotypes. Recent review articles have covered these applications comprehensively.^4,5^ Critical software tools and workflows for executing this complex pipeline are designed and used by bioimage analysts^6^, research software engineers^7^, and others with similar roles.

**Figure 1.**
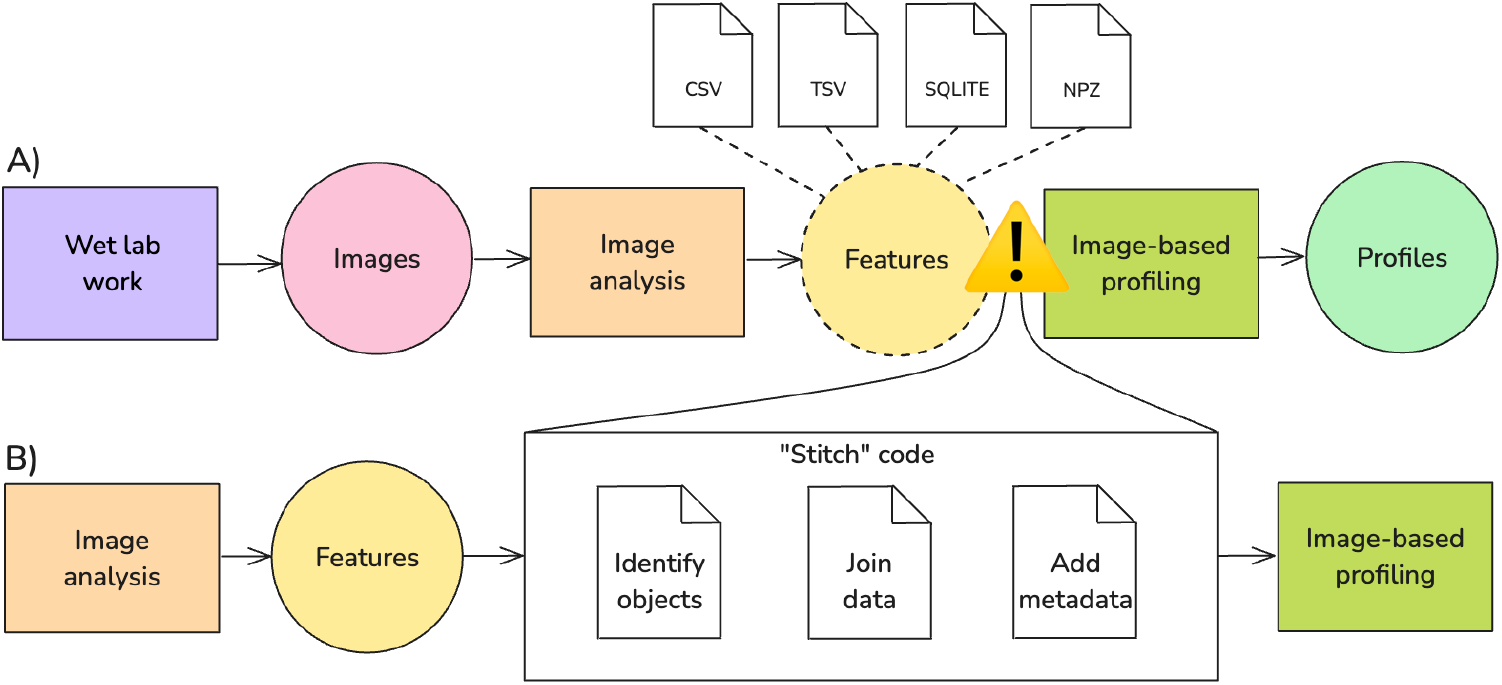
The standard data pipeline or bioinformatics workflow(s) to analyze high-content imaging (HCI) data. **(A)** The HCI pipeline starts with a hypothesis and wet lab work (a process symbolized by a rectangle) that results in the collection of microscopy images (data which are symbolized as a circle). Next, image analysis converts the images into morphology features, which then progress through an image-based profiling pipeline to generate processed profiles for downstream hypothesis testing and biological discovery. Image analysis software produces features that are stored in myriad file formats from several different image analysis tools. The output features from different image analysis tools are inconsistent, posing several challenges (symbolized within the figure as a yield sign). For example, challenges include different naming conventions, numbers of files, and different data values. **(B)** To solve these challenges, image analysts “stitch” code (rewrite existing and bring together previous code iterations) to integrate data prior to image-based profiling steps, which slows progress and may lead to reproducibility concerns.

Image-based profiling data formats and structures are myriad; there is little cohesion on file type, column names, or structure. These data also tend to be “wide”, entailing many thousands of columns, which makes for the high-content nature of the approach. Comma-delimited spreadsheets (e.g., CSV, TSV, etc.) or other text-based data are commonplace and ill-suited to handle large data needs. For example, these formats do not, by default, define explicit data types (e.g., string, float, etc.), which can lead to discrepancies during processing and larger resource requirements to infer data types on-the-fly. Comma-delimited files are also difficult to compress in comparison to database formats (which are optimized to store data), and they are prone to errors such as quoting issues or manually-typed errors, which can be nearly impossible to detect. Other formats, such as SQLite, provide explicit relationships between tables as an advantage, but persist an ambiguous typing system that can result in, for example, strings in floating point columns (leading to similar data inference challenges).

A persistent and under-addressed challenge in image-based profiling is the lack of robust data integration pathways between different image analysis software feature extractors (e.g., CellProfiler^8,9^, DeepProfiler^10^, Molecular Devices IN Carta^11^) and image-based profilers (e.g., Pycytominer^2^) (**Fig. 1B**). While feature extractor software can extract thousands of features from images, they typically require additional processing to become interoperable with downstream profiling. Specifically, these extractors do not provide standardized data or interfaces to support the transformation of features into structures compatible with image-based profiling workflows. As a result, researchers are frequently left to implement custom, *ad hoc* “stitch” code which, for example, borrows and rewrites existing code to join multiple CSVs, harmonize metadata, or reconcile segmentation identifiers in order to take full advantage of image-based profiling software. This process is not only error-prone, but also inhibits reproducibility and scalability across experiments and laboratories by consuming valuable research time. Addressing this integration bottleneck is critical for enabling more automated, scalable, and reproducible HCI pipelines.

## Methods

### CytoTable for feature integration

The increasing complexity and scale of HCI datasets demand computational tools that support scalable, interoperable, and reproducible analyses. The term data integration has been used to describe the technical challenge of combining heterogeneous biological datasets^12^, while more recent work emphasizes data harmonization as the process of reconciling differences in formats, definitions, and measurements across studies.^13^ CytoTable addresses both needs by integrating diverse image-based feature extraction outputs into a unified structure and harmonizing them into consistent, interoperable formats (e.g., Arrow^14^, Parquet^15^, AnnData^16^) suitable for reproducible downstream analysis. We developed CytoTable to bridge this gap as a scalable and reproducible “feature integrator” across many different image-based feature extractors and data formats (**Fig. 2A**). CytoTable facilitates the transformation of raw image-based feature outputs into harmonized, bioinformatics-ready data by resolving complex multi-table relationships and enabling high-performance, single-cell data integration for downstream profiling (**Fig. 2B**).

**Figure 2.**
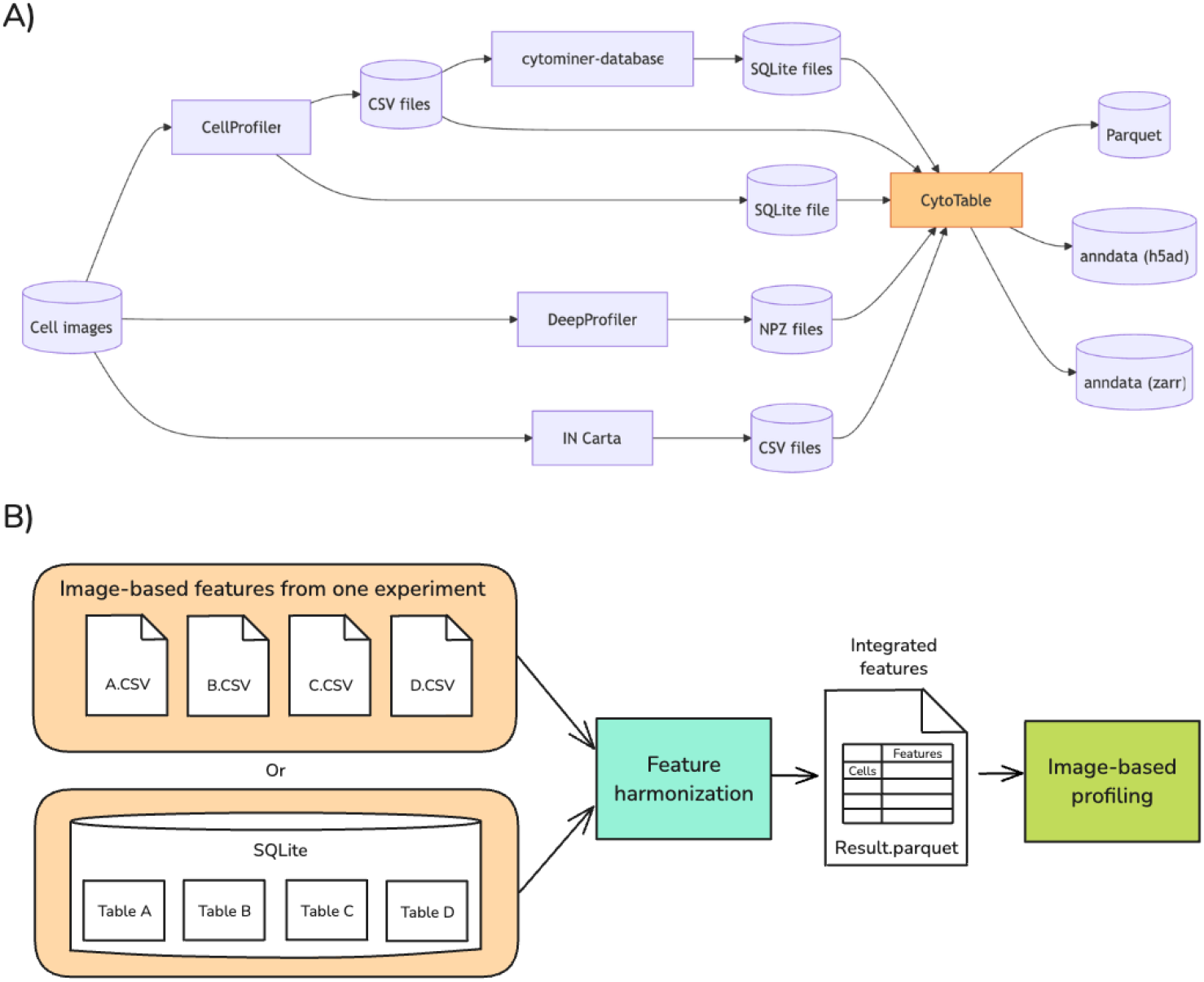
CytoTable solves the important problems of feature harmonization for image-based profiling. **(A)** Cytotable harmonizes a range of data formats commonly used in high-content imaging (HCI) workflows. These data sources originate from different image analysis software tools widely adopted in the HCI community. Each output format is compatible with Pycytominer and other image-based profiling software. **(B)** Through harmonization, CytoTable outputs file formats that integrate into downstream image-based profiling bioinformatics software. Feature harmonization through CytoTable involves consistent column name handling, data typing, and serialization to a number of formats.

CytoTable applies the data integration phase of well-established data-processing and data-mining framework CRISP-DM.^17^ By translating the heterogeneous outputs of individual feature extractors into a single, schema-driven table before any downstream analysis, it removes the tool-specific quirks that usually splinter HCI workflows and limit cross-study reuse. This standardized layer turns CytoTable from a mere Extract, Transform, and Load (ETL) utility into a reproducible, modular backbone on which image-based profiling pipelines can scale confidently and interoperably (**Extended Note 1**).

### CytoTable components

Inspired by modular data engineering principles and the Composable Data Management System Manifesto^18^, we designed CytoTable with a robust framework for managing varied image analysis output files from HCI experiments. By leveraging a unified data representation, optimized query execution, and seamless cross-language interoperability, CytoTable enables researchers to efficiently process and analyze large-scale HCI data.

This conceptual model is grounded in long-term, sustainable software practices^19,20^ designed to promote maintainability alongside adaptability to avoid software collapse.^21^ To meet this evolving landscape, we developed using modular, interoperable components that reflect a commitment to modular and emergent design principles, inspired by the work of the Software Gardening Almanack.^22,23^ The approach not only increases the longevity of CytoTable but also leaves a durable foundation for future software tools and evolving standards in the image-based profiling community.

### Cross-platform in-memory data types with Apache Arrow

At its core, CytoTable employs Apache Arrow^14^ tables through PyArrow (the Pythonic API for Apache Arrow) for an intermediate in-memory representation, ensuring efficient columnar data access and minimizing the time it takes to prepare data within software procedures (**Fig. 3A**). PyArrow tables within CytoTable handle all single-cell features to optimize processing performance and enforce strict typing expectations. Apache Arrow is an open-source, columnar in-memory data format designed to optimize data processing and interoperability across multiple programming languages. This structured approach allows users and maintainers to seamlessly transition between in-memory operations and long-term storage, facilitating reproducible workflows in image-based profiling.

**Figure 3.**
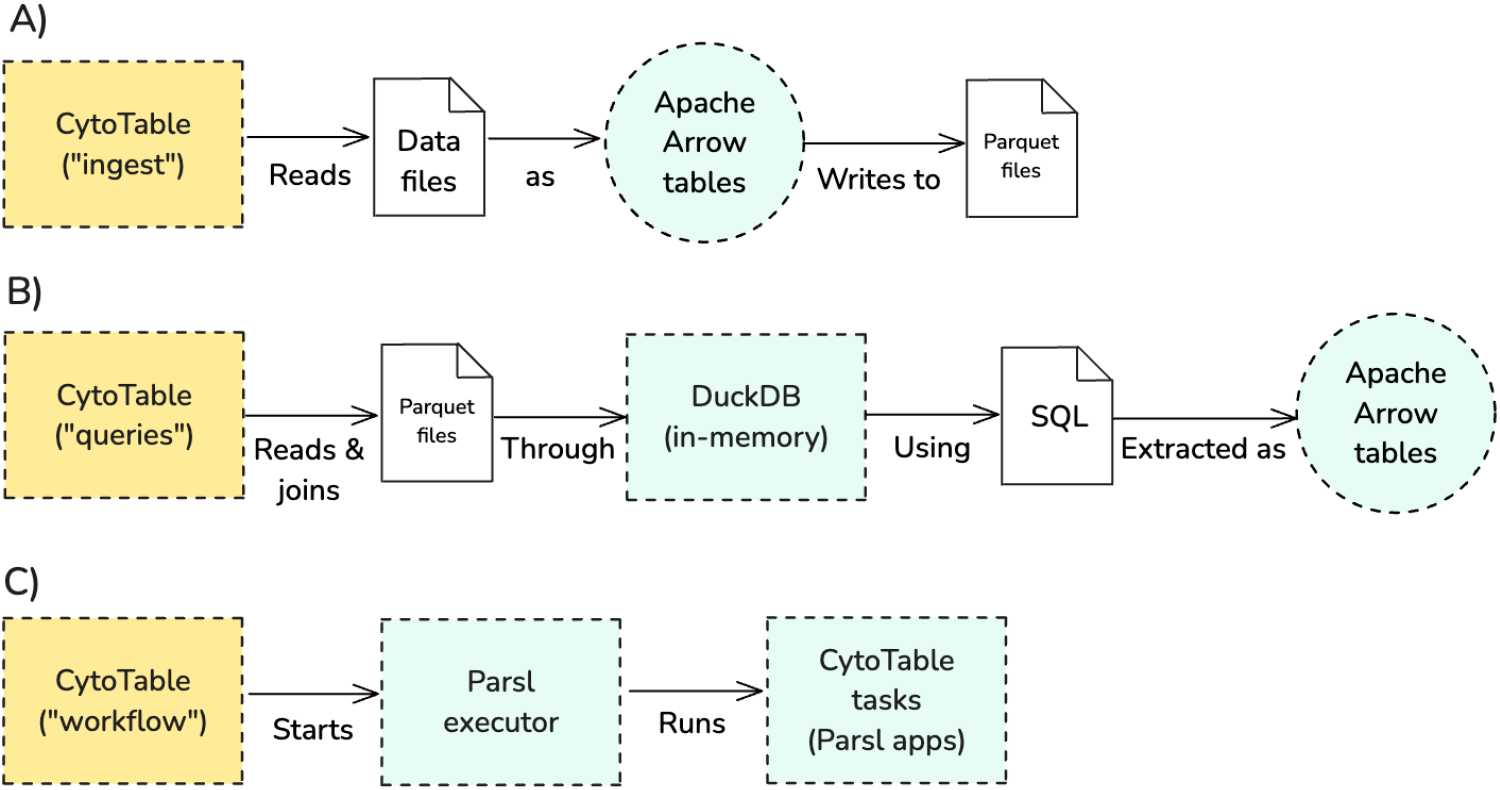
Core CytoTable processes. (**A**) CytoTable ingests serialized data from files as Apache Arrow tables using PyArrow. CytoTable then serializes the data files into Parquet files, retaining strong data type relationships for Arrow integration. (**B**) CytoTable reads and performs data joins through an in-memory DuckDB database as a query engine using SQL to extract Apache Arrow tables. **(C)** CytoTable uses Parsl to orchestrate parallelizable tasks as a data workflow.

### Consistent and performant serialization through Parquet data storage

CytoTable leverages Apache Parquet^15^ as its primary format for persistent data storage, ensuring efficient handling of large-scale single-cell HCI datasets. All intermediate and finalized feature data integrated by CytoTable are exported to Parquet. By adopting Parquet’s columnar storage model, CytoTable significantly reduces disk I/O and improves query performance, particularly for analytical workloads that require scanning and aggregating large amounts of structured data. Its columnar layout is particularly advantageous for analytical queries, as it allows scanning only the necessary columns instead of entire rows, significantly improving query performance. Due to these advantages, Parquet is the foundation of some of the largest data infrastructures in the world, including those used by Google BigQuery^24^, Amazon Athena^25^, Snowflake^26^, and Apache Spark.^27^ Parquet’s compression and encoding capabilities further enhance storage efficiency, making CytoTable well-suited for workflows that involve iterative analysis and large-scale data sharing across computational environments.

### Portable high-performance OLAP with DuckDB and SQL

To enhance cross-platform analytical performance, CytoTable integrates DuckDB^28^, an embedded analytical database designed for high-speed, Structured Query Language (SQL) based queries on datasets (**Fig. 3B**). SQL, a domain-specific language introduced in 1974^29^ for managing data in relational database systems, remains one of the most widely adopted standards for querying and analyzing structured data. Its enduring relevance, portability, and declarative nature made it a natural choice for integration within CytoTable. DuckDB is an embedded analytical database designed for online analytical processing (OLAP), offering high-speed execution of complex queries on structured data. Unlike traditional transactional databases, DuckDB is optimized for read-heavy workloads, leveraging vectorized query execution and automatic optimizations to efficiently handle large datasets. Its ability to execute SQL queries directly within local environments, without requiring a separate database server, makes it an ideal tool for scalable data analysis workflows. Further, DuckDB enables direct querying of Parquet or CSV data stored in cloud object storage (e.g., S3, GCS, Azure Blob), eliminating the need for local downloads and supporting analyses across both local and distributed environments (see **Extended Note 2** for more information on cloud-based data access). CytoTable performs all extraction and join work on single-cell feature data through DuckDB SQL, enabling users to perform complex data transformations and aggregations with minimal computational overhead, scaling even to large-scale datasets (**Extended Note 2**).

### Scalable task-based parallel workflow execution with Parsl

CytoTable leverages Parsl^30^, a parallel programming library for Python, to implement MapReduce-style^31^ workflows and distributed data processing. Parsl enables researchers to define modular, task-based execution pipelines that can scale seamlessly from local machines to high-performance computing (HPC) clusters (**Fig. 3C**). By abstracting task execution into a flexible dependency graph, Parsl facilitates efficient parallel execution of data transformations, distributing SQL queries, filtering operations, and other computational tasks across multiple processing units. This data-driven execution model is rooted in early dataflow semantics introduced by Dennis and Van Horn (1966), where tasks are triggered by the availability of their inputs rather than by a centralized control.^32^ This structure supports both multiprocessing and multithreading by decoupling task scheduling from the limitations of sequential programming.

It is important to understand how Python interacts with CPU cores and threads to realize these dataflow benefits on today’s hardware. Modern computation relies on CPU cores, which are independent compute engines housed within each CPU package. Threads are an operating-system abstraction that the scheduler maps onto those cores. In CPython versions below 3.13, every process contains a Global Interpreter Lock (GIL), a mutex that permits only one thread at a time to execute Python byte-code within that process. CPython versions 3.13 and above enable optional and eventual non-default GIL.^33^ As a result, common and current CPython CPU-bound workloads may achieve true parallelism only with multiprocessing, which launches multiple processes (each with its own GIL) that the operating system (OS) can run concurrently on different cores. Multithreading creates several threads inside a single process; although the OS may place them on separate cores, they still contend for the same GIL, so the approach mainly benefits I/O-bound tasks or native libraries that release the lock. CytoTable supports both multiprocessing and multithreading, orchestrating single-cell data integration through Parsl’s execution engines.

### Accessible, extensible, and interoperable Python interface

CytoTable provides feature integration capabilities to image-based profiling researchers by providing well-documented application programming interfaces (APIs) which can be readily installed as a Python package (e.g., “pip install cytotable”). Once installed, the Python package is importable into Python modules or scripts to flexibly implement feature integration into image-based profiling pipelines where it makes sense. CyoTable takes advantage of features afforded by Python packaging frameworks for automated testing, quality assurance, deployment, document rendering, and other tasks to promote long-term project sustainability in one of the most used programming languages.

CytoTable is highly customizable to accommodate and harmonize multiple input sources To accommodate different data schemas and conventions, CytoTable supports both manually specified configuration arguments and readymade presets, which are included at installation. The configuration arguments enable flexibility when it comes to data ingestion, processing, and exports. Configuration presets (e.g., “cellprofiler_csv” or “in-carta”) allow users to apply prepared configuration arguments for CytoTable in common data format expectations without handcrafting specific SQL queries or other customizations (**Table 1**). When presets do not suffice, users can override specific options directly via arguments (allowing for partial use of presets). Each preset is versioned with the image analysis software, and they are included in software tests to ensure deterministic and accurate results. A special parameter, “joins”, within these presets helps document the SQL-based relationship between tables, which are extracted from image analysis software. These presets are helpful in using CytoTable for known data sources or adjusting to meet new configurations.

**Table 1.** CytoTable provides quick configuration options through the use of “presets” which are supplied at runtime. Presets allow users to take advantage of common data formats output by image analysis software to save time and reach data output sooner. Presets may be completely or partially overridden with individual CytoTable argument values for customized implementations as needed.

| Software | Software output format | CytoTable preset |
| --- | --- | --- |
| CellProfiler | CSV | cellprofiler_csv |
| CellProfiler | SQLite | cellprofiler_sqlite |
| DeepProfiler | NPZ | deepprofiler |
| IN Carta | CSV | in-carta |
| cytominer-database | SQLite | cell-health-cellprofiler-to-cytominer-database |

CytoTable supports image-based profiling outputs from CellProfiler, which is an open-source image analysis software.^8,9^ It is widely used for cell segmentation, image quality control, and morphology feature extraction. It is often used in high-throughput screening, which generates many terabytes of data and demands high compute costs.^34,35^ CellProfiler produces image analysis outputs in many different forms, including CSV and SQLite data (**Table 1**). CellProfiler exports in CSV or SQLite formats, based respectively on the *ExportToSpreadsheet* or *ExportToDatabase* modules. CytoTable accommodates either output format with the existing presets: “cellprofiler_csv” and “cellprofiler_sqlite”. Additionally, cytominer-database is a deprecated tool that cleans and processes CellProfiler output CSV files. We created a preset specially designed for this data called: “cell-health-cellprofiler-to-cytominer-database”. Because of its deprecation, we expect that this preset will only support legacy CellProfiler output data which scientists may want to reanalyze.

CytoTable also supports image-based profiling outputs from DeepProfiler, which is an open-source deep learning pipeline developed by the Cytomining community to generate morphological profiles directly from images using pre-trained deep learning models.^10^ DeepProfiler outputs single-cell feature matrices in .npz format, which is a compressed NumPy archive suitable for efficient handling of high-dimensional data arrays. CytoTable provides the “deepprofiler” preset to harmonize these data.

Lastly, CytoTable supports image-based profiling outputs from IN Carta, which is a commercial image analysis software platform developed by Molecular Devices^11^ to provide tools for cell segmentation, tracking, and feature extraction. IN Carta exports processed image-based features as CSV files. These files contain per-object or per-image measurements suitable for morphological profiling and downstream analytics. CytoTable provides the “in-carta” preset for use with this format. Importantly, customizing presets is possible through overrides and customization. We expect that as other image analysis feature data formats are developed that we (or other community members) can quickly develop new customized presets.

### CytoTable benchmarking with other platforms and tools

We benchmarked join operations and memory usage between CytoTable and Pycytominer on realistic single-cell profiling tasks. Pycytominer is a bioinformatics software for processing image-based profiling data, and it contains a utility class called SingleCells. This class, which is much more limited in scope than CytoTable, harmonizes CellProfiler output data prior to bioinformatics processing. We ran each benchmark six times per file type to quantify performance fluctuations.^36^ We benchmarked using a Linux desktop computer with 16 CPU cores and 64 GB of RAM. Because of limited functionality in Pycytominer, benchmarking comparisons between Pycytominer and CytoTable could only be directly performed using SQLite datasets. Therefore, our CSV dataset benchmarking used the Pandas^37^ Python library rather than Pycytominer directly. All CytoTable comparisons involved two different Parsl execution engines: HighThroughputExecutor and ThreadPoolExecutor, labeled “multiprocess” and “multithread” respectively. We used Parsl’s MonitoringHub^38^ to profile memory for the HighThroughputExecutor because it involves specialized process spawning, which otherwise may result in inaccurate memory profiles. We performed all other memory profiling using psutil^39^, a commonly used and widely accepted tool within the Python community. We strove to provide an honest, best effort benchmarking of these technologies.^40^

## Results

### CytoTable comparison benchmarks

As image-based profiling workflows grow in scale and complexity, efficient data handling becomes increasingly important. We developed CytoTable in response to practical limitations encountered in earlier workflows which revolve around in-memory operations using Pandas^37^ DataFrames. While Pandas provides a flexible interface for data manipulation, its performance can degrade significantly when handling large-scale datasets, particularly during join operations, where multiple tables must be merged based on shared keys. These joins often require entire tables to be loaded and sometimes duplicated into memory, resulting in substantial resource consumption and potential scalability bottlenecks. A primary motivation for CytoTable was to reduce these memory requirements by leveraging more efficient data representations and execution strategies, enabling scalable data processing without compromising performance. We also developed CytoTable as a harmonization tool, which could accept multiple different image-analysis outputs. Pycytominer does not have this functionality, and we therefore wanted CytoTable to support newcomers (beyond CellProfiler users) into the Cytomining image-based profiling community.

When processing large CSVs, CytoTable uses several gigabytes less peak RAM than Pandas (**Fig. 4A**), though Pandas still completes a few seconds faster (**Fig. 4B**). For SQLite inputs, CytoTable again requires far less memory than Pycytominer (**Fig. 4C**) and scales linearly, whereas Pycytominer’s run time rises exponentially once file size exceeds 230 MB (**Fig. 4D**). Within CytoTable itself, the default multiprocessing mode delivers the smallest memory footprint, while multithreading offers the shortest wall-clock time. Overall, Pandas is preferable for small CSV files, and Pycytominer’s SingleCells.merge_single_cells method is preferable for small SQLite files, but CytoTable scales more gracefully as datasets grow, especially for SQLite files, providing a better speed-versus-memory trade-off. Ongoing work aims to further accelerate CSV parsing without sacrificing CytoTable’s memory efficiency.

**Figure 4.**
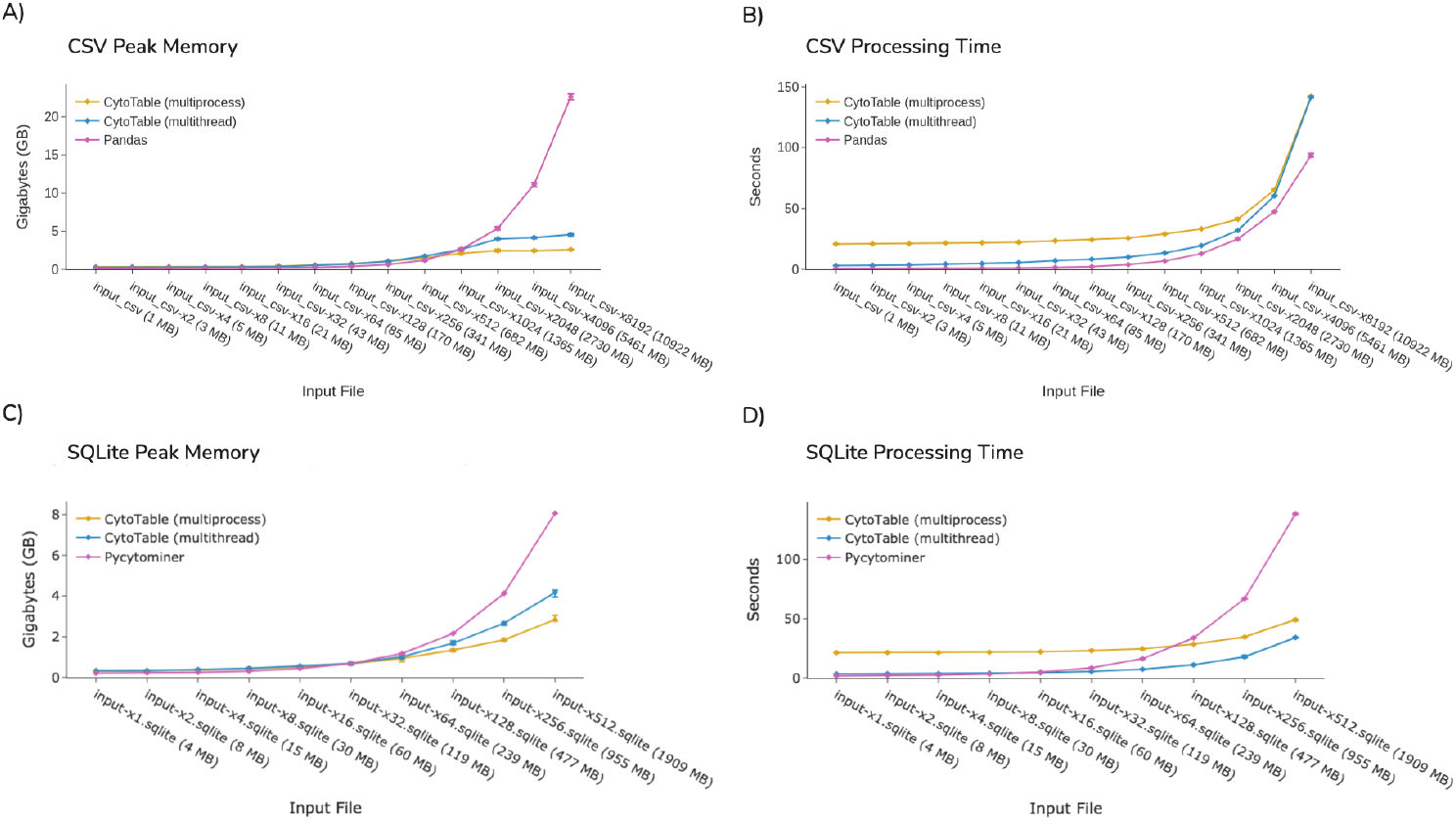
Benchmarking CytoTable and Pycytominer processing with increasingly large input data based on the NF1 Schwann Cell Project (simulating larger data sizes with duplicated records). **(A)** CytoTable outperforms Pandas for CSV-based memory performance. **(B)** CytoTable outperforms Pandas in processing time for CSVs. **(C)** CytoTable outperforms Pycytominer for SQLite memory performance. **(D)**. CytoTable outperforms Pycytominer in processing time for SQLite files.

We chose to develop CytoTable using Apache Arrow, striking a balance between performance and flexibility. While tools like Pandas offer a user-friendly API and are widely adopted in data science, they can be inefficient when scaling to large datasets due to their reliance on dense in-memory structures. PyArrow serves as a common foundation across these ecosystems, offering a language-agnostic, columnar memory format that minimizes serialization costs and supports zero-copy data exchange. CytoTable adopts PyArrow Tables as its internal data representation to enable interoperability and standardize data types between file format and in-memory use. We observed that Apache Arrow tables through PyArrow or Polars^41^ (another Arrow-compatible Python package) use roughly half the memory of Pandas DataFrames (**Fig. 5A**).

**Figure 5.**
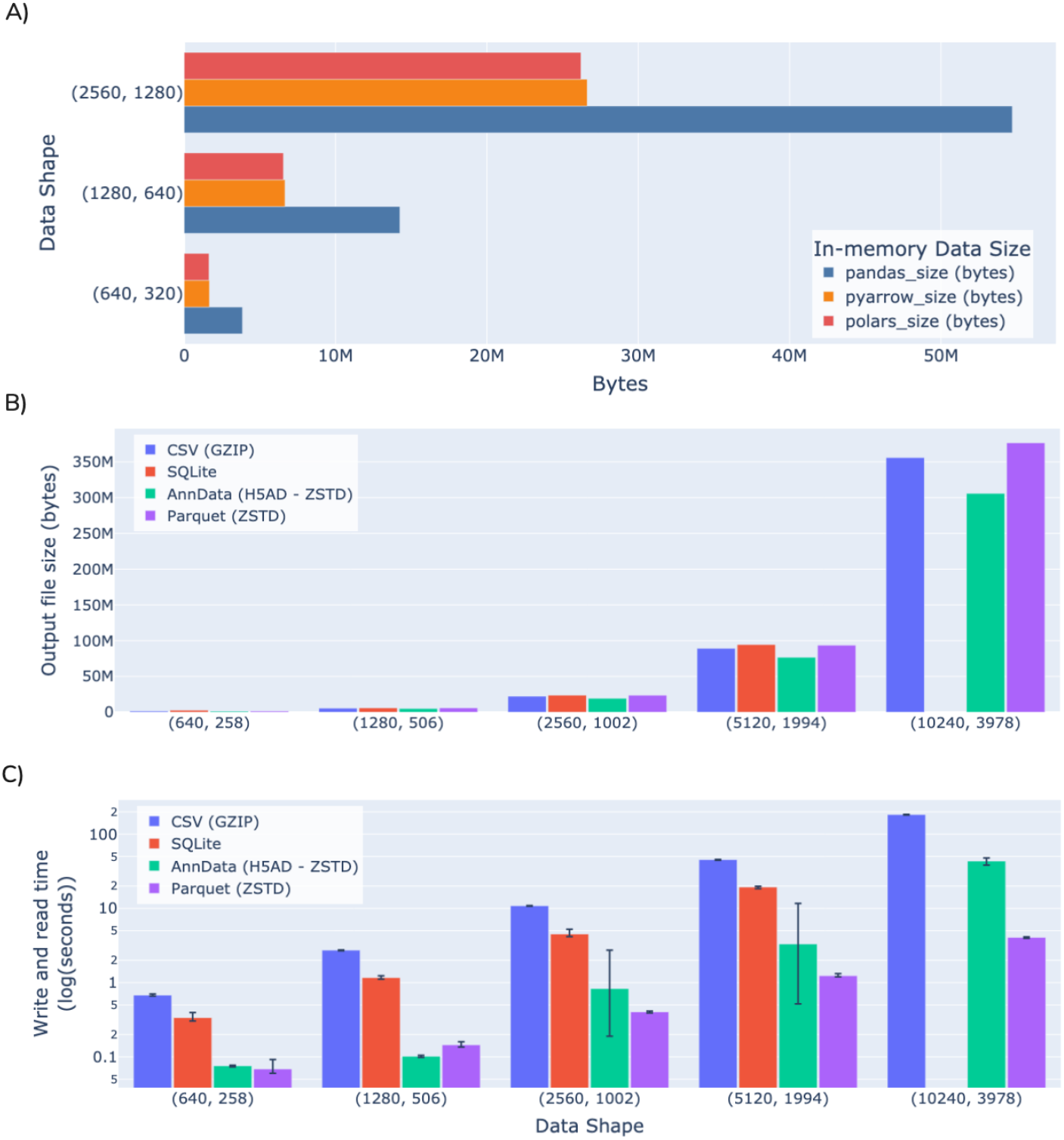
Benchmarking storage and file format performance. (**A**) Loading data through PyArrow tables uses almost half as much memory as Pandas DataFrames. We also compare Polars DataFrames which leverage Arrow constructs to help contextualize how Arrow uses memory in different packages. (**B**) Comparing output file sizes. AnnData achieves lower data storage when compared to CSV, SQLite and Parquet. Parquet is equivalent in data storage size or sometimes larger than CSV or SQLite files. (**C**) Comparing write and read performance. Parquet consistently achieves better write and read time duration performance when compared with AnnData, SQLite, and CSV. Note that SQLite is incompatible for the largest data shape, given the default maximum column length of 2000. Error bars represent the range of six independent runs.

Choosing the appropriate file format is critical for working with high-throughput image-based profiling data. CSV, though commonly used, lacks built-in support for types, compression, or metadata, which limits its suitability for large-scale, structured data. SQLite improves on this by enabling queryable, relational storage in a single file, but it is not optimized for analytical workloads (table columns are by default limited to a maximum of 2000 columns) and has limited parallel I/O capabilities. In contrast, CytoTable uses Apache Parquet by default for its persistent storage format. Parquet’s columnar layout and native compression make it highly efficient for storing and retrieving subsets of large datasets, especially in scenarios involving filtering or aggregation. Its compatibility with modern data systems and ability to handle high-throughput data access patterns make it a natural choice for image-based profiling pipelines. We observed that Parquet may have roughly the same data storage size properties as SQLite or CSV files (**Fig. 5B**) but outperforms both formats when it comes to read (**Fig. 5C**) and write time durations (**Fig. 5C**).

Beyond CSV and SQLite, AnnData (commonly used in the single-cell transcriptomics community) provides a rich container format for storing both data matrices and associated metadata. In benchmarking we found that AnnData results in 17.5% smaller files on average than CSV, SQLite, or Parquet files (**Fig. 5B, Fig. 6**). Parquet consistently outperforms AnnData in read and write times (up to 90% better, 34% average), even when both rely on identical compression algorithms (**Fig. 5C**). In addition, Parquet supports partial writes and updates through multi-file dataset structures, which allow efficient appending or partitioning of large-scale data. Equivalent functionality in AnnData is currently only available through experimental projects such as AnnCollection. These practical advantages make Parquet a more scalable and flexible option for handling high-throughput image-based profiling data within CytoTable.

**Figure 6.**
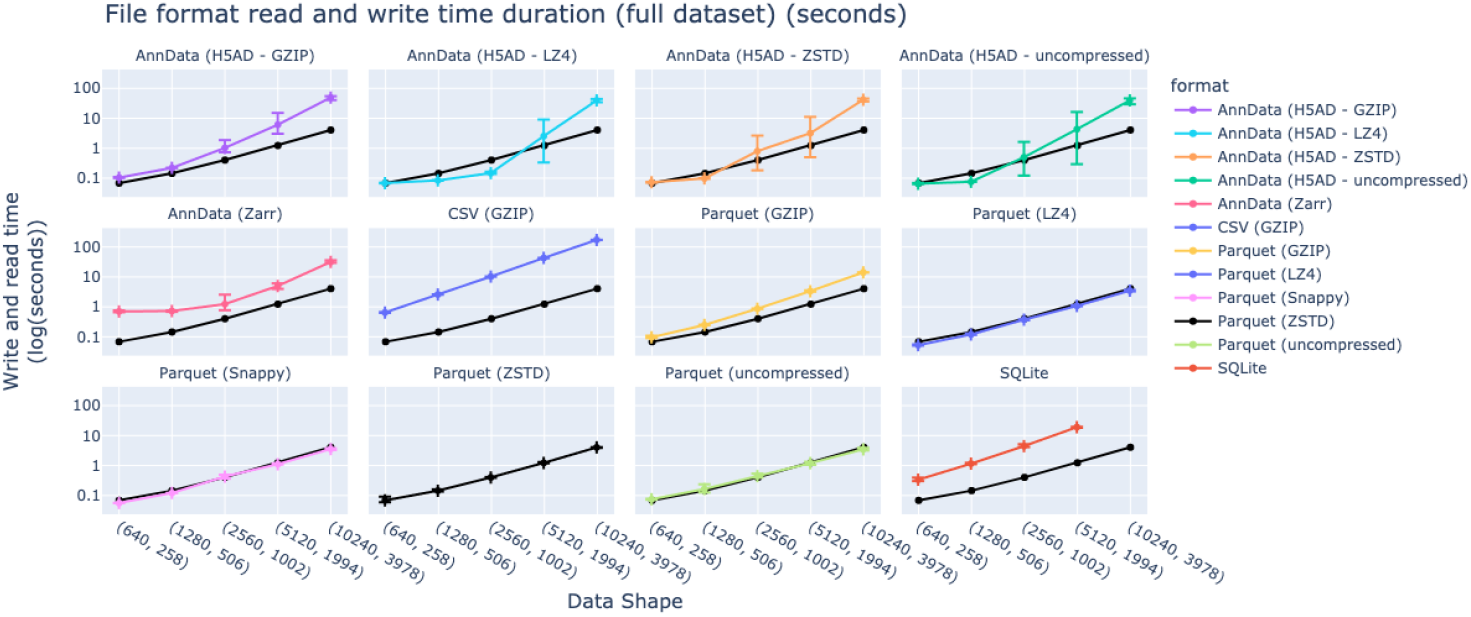
Benchmarking read and write time across data formats with varying compression options. Parquet has consistently better time performance for full dataset reads and writes (including through various compression algorithms) when compared to CSV, SQLite, and several AnnData formats. Note that SQLite is incompatible for the largest data shape, given the default maximum column length of 2000. Error bars represent the range of six independent runs.

## Applied use cases of CytoTable

To demonstrate CytoTable’s practical utility across diverse experimental contexts, we highlight several applied use cases where it has already been integrated into real-world HCI workflows (**Table 2**). These examples span both small- and large-scale datasets, showcasing CytoTable’s ability to standardize image-analysis outputs, support quality control, and streamline downstream analyses to enable reproducible, scalable data integration across a range of applications. These examples contextualize real-world projects with the benchmarks in the previous sections, illustrating that benchmarking expectations are mappable to applied use cases. Specifically, Figs. 4 and 5 leveraged data from a single SQLite file from the NF1 Schwann Cell Project ranging from 242 to 123,904 single cells. Fig. 6 used simulated datasets where each row represents a single cell (ranging from 640 to 10,240 simulated single cells).

**Table 2.**
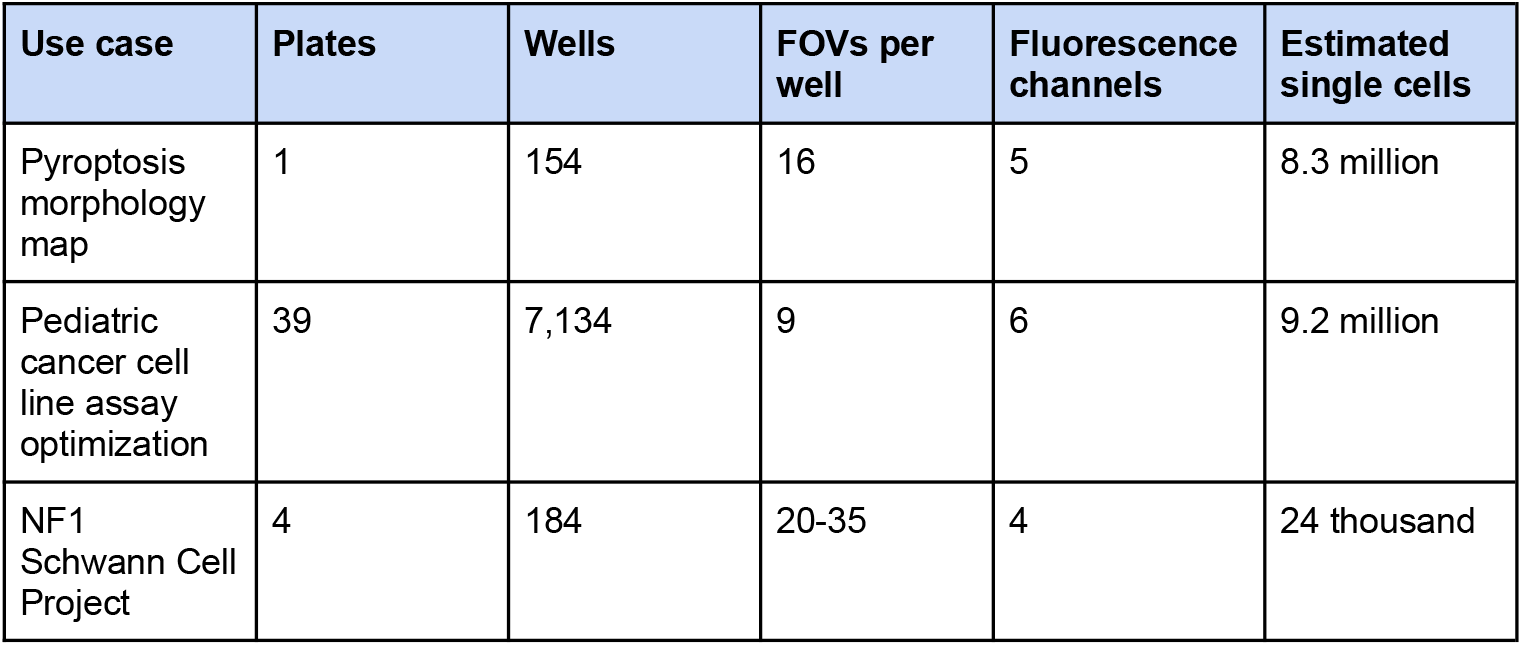
CytoTable has been used in various research projects with varying data sizes to achieve image-based profiling harmonization. Each project used customized versions of the CytoTable preset “cellprofiler_sqlite_pycytominer”.

### Pyroptosis morphology map

Building on similar single-cell profiling workflows, “*A morphology and secretome map of pyroptosis*”^42^ employed CytoTable to process data from over eight million peripheral blood mononuclear cells (PBMCs) subjected to various treatments to induce specific forms of cell death. Here, CytoTable was integrated into an image-based profiling pipeline alongside CellProfiler, facilitating the extraction and harmonization of nearly 3,000 features per cell. The formatted data enabled multimodal analyses linking morphology to inflammatory secretome responses, providing insights into mechanisms of programmed cell death. This effort involved one plate with 154 wells, 16 FOVs and around 8.3 million single cells (**Table 2**).

### Pediatric cancer assay optimization

In an effort focused on optimizing experimental conditions for the Cell Painting assay applied across pediatric cancer cell lines, CytoTable was used to process morphological data.^43^ Following segmentation and feature extraction by CellProfiler, CytoTable structured the single-cell profiles to support downstream processing with coSMicQC^44^ and Pycytominer^2^. This formatting step helped evaluate how factors like seeding density and assay timing influenced segmentation quality and morphological consistency, contributing to improved reproducibility in large-scale profiling workflows. This effort involved 24 plates with varied well counts and six total channels imaged (**Table 2**).

### NF1 Schwann Cell Project

In the study “*High-content microscopy and machine learning characterize a cell morphology signature of NF1 genotype in Schwann cells*,”^45^ researchers applied a modified Cell Painting assay to profile single-cell morphology in Schwann cells across different *NF1* genotypes. CytoTable was used to curate and format the high-dimensional segmentation outputs, enabling downstream integration with tools such as Pycytominer and coSMicQC. The resulting dataset of over 20,000 single-cell profiles supported quality control and statistical analyses, ultimately revealing subtle morphological signatures associated with *NF1* genotype. This effort involved four plates with varied well counts and four channels (**Table 2**).

### Cytomining ecosystem project integration

Beyond individual studies, CytoTable plays a central role within the Cytomining ecosystem by providing standardized outputs that feed directly into downstream tools. Cytomining is a GitHub organization and community that develops and hosts image-based profiling bioinformatics software like Pycytominer.^2^ Pycytominer takes CytoTable data as input for image-based profiling workflows; coSMicQC uses CytoTable data to define thresholds for reliable quality control; and CytoDataFrame extends CytoTable output into interactive, in-memory exploration of single-cell profiles alongside images and segmentation masks. Together, these integrations demonstrate how CytoTable serves as a connective layer across Cytomining projects, enabling reproducible pipelines that span data harmonization, quality control, and exploratory analysis within the broader ecosystem of image-based profiling.

## Discussion

CytoTable is a core component of the Cytomining ecosystem, which is a vibrant open-source community focused on developing interoperable tools for HCI and high-throughput image-based profiling. The data outputs from CytoTable support streamlined input into downstream software such as coSMicQC^44^, which performs rigorous quality control on single-cell segmentation results, and Pycytominer^2^, a flexible framework for processing image-based features into formats compatible with statistical and machine learning workflows. While each of these tools offers powerful capabilities on their own, they depend on a consistent, scalable data infrastructure to effectively bridge the gap between raw image-derived measurements from image analysis tools and actionable biological insights. By delivering a standardized, memory-efficient, and SQL-compatible format for single-cell image-based profiling, CytoTable serves as a key integrative layer in this ecosystem.

As the image-based profiling field has matured through practical use and iterative development, several technical challenges and design considerations have emerged.^3^ These considerations include in-memory and serialized data types, consistent floating point numeric precision capture within in-memory data types, and memory allocator effectiveness combined with data types such as Apache Arrow (**Extended Note 4**). These experiences have informed how CytoTable handles data format variability, numerical precision, and memory management across different execution environments. CytoTable solves these challenges for the image-based profiling field.

In the spirit of *panta rhei*—that everything flows and nothing stays the same—we acknowledge that the HCI and image-based profiling landscapes are constantly shifting. New tools, formats, and workflows continue to emerge, and any software aiming to support this community must evolve alongside it. CytoTable is built with this adaptability in mind: not as a rigid framework, but as a modular system designed to meet the changing needs of a diversity of bioimage informatics tasks. Looking ahead, we outline several key areas where CytoTable will continue to grow in response to this evolving ecosystem. CytoTable will continue to evolve as part of the broader Cytomining ecosystem, benefiting from shared standards, complementary tools, and an active open-source community. Iterative enhancements will focus on performance and scalability while maintaining strong interoperability with formats and workflows emerging across bioimaging and scverse^46^ projects. By anchoring development within this collaborative ecosystem, CytoTable can adapt quickly to community needs, new image analysis technologies, sustain reproducibility through shared testing and release practices, and provide a reliable foundation for advancing image-based profiling.

CytoTable is not without limitations. A current limitation of CytoTable is that its harmonization approach depends on morphology feature schema conventions and file formats that continue to evolve. As standards shift, sustained interoperability will require ongoing alignment and adaptation. While the open-source, ecosystem-driven model provides a pathway for addressing these challenges, it also means that reproducibility may be temporarily constrained if upstream tools or formats change faster than CytoTable can incorporate updates.

A central challenge in image-based profiling is the diversity of image analysis formats, schemas, and workflows that make integration difficult.^47^ CytoTable contributes toward a shared data model by harmonizing heterogeneous inputs into consistent, analysis-ready outputs, while remaining flexible enough to accommodate emerging tools and standards. Continued progress in this direction will reduce duplication, lower barriers to interoperability, and support the long-term goal of a collaboratively developed model for single-cell morphology data across multiple organizations (such as that of scverse).

### Future development: Expanding input sources

While CytoTable currently provides robust support for standard image-based profiling outputs, future development will expand its compatibility with a broader range of image analysis input types through additional presets and custom configurations. Currently tailored to formats such as CellProfiler’s CSV and SQLite exports, upcoming versions aim to natively support additional tools and pipelines in the image-based profiling ecosystem. This expansion will improve interoperability across profiling tools and reduce the need for custom preprocessing scripts, making CytoTable a more flexible entry point into standardized single-cell morphology analysis workflows.

### Future development: Enhancing output format options

Emerging formats such as Lance^48^, Nimble^49^ (initially called “Alpha”), and Vortex^50^ offer new opportunities for scalable data storage, streaming access, and versioned datasets optimized for analytical performance and cloud-native environments. Integrating these formats will provide users with greater flexibility in managing large-scale single-cell datasets, particularly in settings that demand fast I/O, partial reads, or remote data access. Lance in particular includes array-value capabilities which are a gap of Parquet and could be leveraged to great effect when it comes to features, image data (as arrays), or shape vectors (of image objects). It also is optimized for vector search which could offer benefits to profiling efforts. AnnData^16^ is another potential output format option to integrate with other tooling in the scverse^47^.

### Future development: Integrating with CytoDataFrame

Finally, we plan a deeper integration with CytoDataFrame^51^, an in-memory data abstraction for single-cell image-based profiling. We designed CytoDataFrame to couple cell measurements with associated metadata, segmentation masks, and raw images in memory, supporting interactive analysis and visualization in environments like Jupyter. CytoTable, in contrast, is focused on feature harmonization and serialization into file formats. Future work will explore bi-directional compatibility between CytoTable and CytoDataFrame (e.g., enabling .to_cytotable() and .from_cytotable() methods) to bridge persistent storage and live, human-in-the-loop exploration that is required for iterative development and hypothesis generation.

## Resource Availability

### Lead contact

Requests for further information and resources should be directed to and will be fulfilled by the lead contact, Gregory P. Way.

## Materials availability

No new materials were generated in this study.

## Data and code availability

- CytoTable is an open-source project and its source code can be viewed and downloaded from https://github.com/cytomining/cytotable.
- CytoTable’s installation and usage documentation is available at https://cytomining.github.io/CytoTable/.
- A brief tutorial on how to use CytoTable is available at https://cytomining.github.io/CytoTable/tutorial.html.
- A notebook based example notebook showing CytoTable in use is available at: https://cytomining.github.io/CytoTable/examples/cytotable_mise_en_place.html.
- A notebook based example notebook showing CytoTable in use with cloud data is available at: https://cytomining.github.io/CytoTable/examples/cytotable_from_the_cloud.html.
- The repository containing the code used to conduct benchmarking and generate results is available at https://github.com/cytomining/CytoTable-benchmarks.

## Declaration of interests

KIB is an employee of Seqera Labs S.L. The authors declare no other competing interests.

## Declaration of generative AI and AI-assisted technologies in the writing process

During the preparation of this work the authors used OpenAI’s ChatGPT in order to improve the readability and language of the manuscript. After using this tool/service, the authors reviewed and edited the content as needed and take full responsibility for the content of the published article.

## Acknowledgments

JT, DB, and GPW were supported in part by Alex’s Lemonade Stand Foundation ‘A’ Award and Tap Cancer Out to GPW (Grant # 23–28306). GPW and ES were supported in part by an American Heart Association Collaborative Sciences Award (24CSA1255857) to GPW. Special thanks goes to the following for their help in contributing to CytoTable design and development or related work.

- Way Lab (https://www.waysciencelab.com/): Cameron Mattson
- Broad Institute (https://www.broadinstitute.org/): Shantanu Singh, Beth Cimini, Sam Chen
- Yale School of Medicine (https://medicine.yale.edu/): Samir Amin
- University of Wisconsin-Madison Morgridge Institute for Research (https://morgridge.org/): Juan C. Caicedo, Nikita Moshkov
- Sanford Burnham Prebys (https://sbpdiscovery.org/): Alexandre Colas, Aashna Lamba

## Author contributions

Conceptualization, DB and GPW; data curation, DB, JT, and ES; formal analysis, DB and JT; funding acquisition, GPW; investigation, DB, JT, ES, MJL, KIB, and GPW; methodology, DB, JT, ES, MJL, KIB, FA, and GPW; project administration, DB and GPW; resources, VR and GPW; software, DB, JT, ES, MJL, KIB, FA, and GPW; supervision, DB and GPW; validation, DB and FA; visualization, DB; writing – original draft, DB and GPW; writing – review & editing, DB, JT, ES, MJL, KIB, VR, FA, and GPW.

## Notes

https://github.com/cytomining/CytoTable

### Summary of Updates

Minor clarifications of text, adding table linking real-world use cases to performance benchmarks.

