## Extended Notes for "Scalable data harmonization for single-cell image-based profiling with CytoTable"

### Extended Note 1 - CytoTable Framework Philosophy

By framing high-content imaging (HCI) explicitly within data mining methodologies, we can better modularize image-based profiling steps, ensure interoperability, and support reproducibility. This modular perspective not only appreciates the strengths of existing tools but also encourages broader collaboration across the open-source ecosystem, ensuring that HCI evolves as a cohesive and community-driven scientific practice.

Within the CRISP-DM (Cross-Industry Standard Process for Data Mining) framework, data integration is recognized as a foundational stage in preparing datasets for analysis.[19] Positioned between data understanding and modeling, this phase ensures that disparate sources, which are often heterogeneous in format and schema, are transformed into a coherent structure suitable for downstream interpretation and modeling. In the context of HCI, where multiple outputs from segmentation pipelines, metadata annotations, and image-derived features must be reconciled, robust data integration is especially critical. More broadly, the lines between data mining and data science blur: while data science emphasizes exploration, hypothesis generation, and flexible analytical tooling, data mining traditionally focuses on producing structured, repeatable outputs such as predictions, classifications, or segmentations through well-defined pipelines.[52] Effective data integration lies at the heart of both, serving as the bridge between raw inputs and actionable insights. As such, tools like CytoTable play an essential role in operationalizing this step, enabling both scientific exploration and reliable downstream application.

### Extended Note 2 - CytoTable data methods

CytoTable is a modular, scalable data processing library designed for transforming and managing image-derived, high-content, single-cell data. To accommodate diverse workflows and ensure flexibility across computing environments, CytoTable integrates task-based execution via Parsl. Below, we describe the architecture of CytoTable workflows and how they integrate cloud-native data access, efficient in-memory operations, and flexible configuration mechanisms.

#### CytoTable pagination

Working with large datasets can pose significant challenges, particularly in environments with constrained memory resources that cannot accommodate entire tables in memory simultaneously. For instance, when loading a full table with Pandas[37], the system must allocate enough memory to hold the entire dataset. If the system doesn't have enough memory, Python may crash or encounter out-of-memory errors. CytoTable addresses this limitation through data pagination, a method that processes data in smaller, manageable subsets (often referred to as "chunks") to avoid memory bottlenecks (**Extended Note 2, Fig. 1**). CytoTable controls pagination behavior by two parameters: "chunk_size", which defines the maximum number of rows per page, and "page_keys", which specify the column(s) used to partition the data. Using these settings, CytoTable dynamically generates data pages and exports each as an individual

Parquet file. Together, these files form a larger Parquet “dataset,” enabling scalable and memory-efficient table processing.

<table>
<tr><th>Original</th><th>Changes</th></tr>
<tr><td>“Data source”
<table>
<tr><th>Col_A</th><th>Col_B</th><th>Col_C</th></tr>
<tr><td>1</td><td>a</td><td>0.01</td></tr>
<tr><td>2</td><td>b</td><td>0.02</td></tr>
</table>
</td><td>Page or “Chunk” 1
<table>
<tr><th>Col_A</th><th>Col_B</th><th>Col_C</th></tr>
<tr><td>1</td><td>a</td><td>0.01</td></tr>
</table>
Page or “Chunk” 2
<table>
<tr><th>Col_A</th><th>Col_B</th><th>Col_C</th></tr>
<tr><td>2</td><td>b</td><td>0.02</td></tr>
</table>
</td></tr>
</table>

**Extended Note 2, Figure 1.** *CytoTable paginates data sources into pages or “chunks”, which are sets of records from the data source, to help reduce memory strain and increase scalable performance.*

#### CytoTable data concatenation and joins

CytoTable supports two core methods for reconstructing and integrating data across paginated or distributed sources: concatenation and joins. Concatenation refers to the vertical stacking of multiple data “chunks” that share identical column schemas, effectively reassembling a full dataset from previously partitioned subsets (**Extended Note 2, Figure 2**). This process reverses the pagination operation and is implemented using a sequential Parquet writer to combine multiple Parquet files into a unified dataset without loading the entirety of the concatenated dataset.

<table>
<tr><th>Original</th><th>Changes</th></tr>
<tr><td>Page or “Chunk" 1
<table>
<tr><th>Col_A</th><th>Col_B</th><th>Col_C</th></tr>
<tr><td>1</td><td>a</td><td>0.01</td></tr>
</table>
Page or “Chunk" 2
<table>
<tr><th>Col_A</th><th>Col_B</th><th>Col_C</th></tr>
<tr><td>2</td><td>b</td><td>0.02</td></tr>
</table>
</td><td>“Concatenated data”
<table>
<tr><th>Col_A</th><th>Col_B</th><th>Col_C</th></tr>
<tr><td>1</td><td>a</td><td>0.01</td></tr>
<tr><td>2</td><td>b</td><td>0.02</td></tr>
</table>
</td></tr>
</table>

**Extended Note 2, Figure 2**. *Concatenation of pages of tabular data with the same columns into singular tables.*

In contrast, joins enable horizontal merging of datasets with overlapping key columns but differing auxiliary data (e.g., metadata)(**Extended Note 2, Figure 3**). These join operations are expressed using DuckDB SQL syntax within CytoTable's `convert()` function, allowing users to declaratively specify SQL-style semantics such as LEFT JOIN or INNER JOIN. While some tools use the term "merge" for similar operations (e.g., `pandas.DataFrame.merge()`), CytoTable explicitly adopts SQL-style "join" terminology to maintain clarity and consistency with database engines like DuckDB, which it leverages under the hood. We use this methodology because data joins through the Pythonic API of DataFrame libraries such as Pandas, Polars, Dask, or Modin all differ in parameter or operation making it otherwise more difficult to discern consistency.

**Original**

"Table 1" (notice **Col_C**)

| Col_A | Col_B | Col_C |
|---|---|---|
| 1 | a | 0.01 |

"Table 2" (notice **Col_Z**)

| Col_A | Col_B | Col_Z |
|---|---|---|
| 1 | a | 2024-01-01 |

**Changes**

"Joined data" (Table 1 left-joined with Table 2)

| Col_A | Col_B | Col_C | Col_Z |
|---|---|---|---|
| 1 | a | 0.01 | 2024-01-01 |

**Join Specification in SQL**

```sql
SELECT *
FROM Table_1
LEFT JOIN Table_2 ON
Table_1.Col_A = Table_2.Col_A;
```

**Extended Note 2, Figure 3**. *Data joins allow for data table sources with joinable keys but disparate columns to be brought together as a new dataset of mixed columns based on the specification of the join (for example, using SQL).*

Relational joins are central to many bioimaging workflows, especially when combining metadata with single-cell measurements. While Pandas' merge() function is commonly used, it requires full in-memory loading of datasets and can be resource-intensive. SQLite supports basic joins, but lacks the performance optimizations necessary for large-scale analytical queries. CytoTable instead relies on DuckDB, an embedded analytical SQL engine designed for efficient local execution on structured data. DuckDB executes joins using a vectorized query engine and operates directly on Arrow and Parquet data, eliminating unnecessary serialization and allowing for performant, scalable data integration. This makes it especially well-suited for workflows that involve complex filtering, aggregation, or cross-table joins in resource-constrained environments. Through benchmarking we observed that DuckDB outperforms Pandas for time duration and memory consumption with larger files, whereas Pandas outperforms DuckDB with smaller data (**Extended Note 2, Figure 4**).

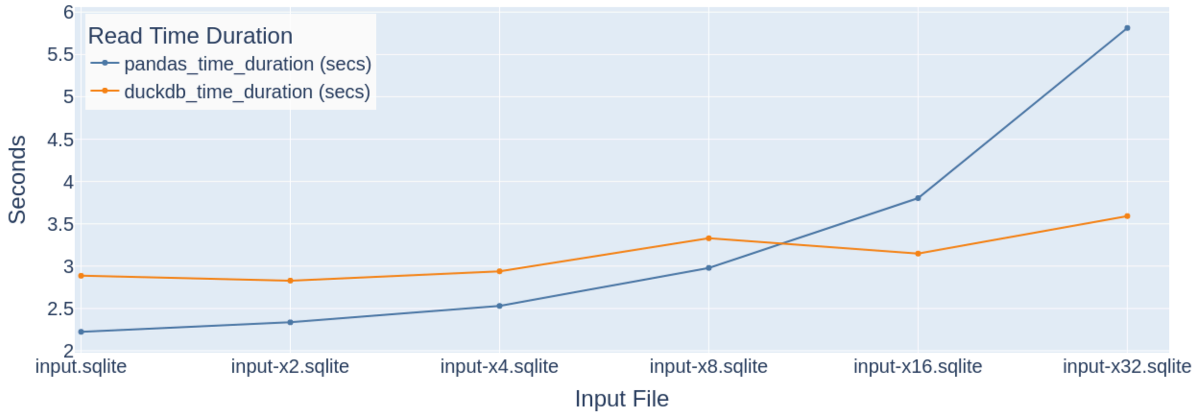


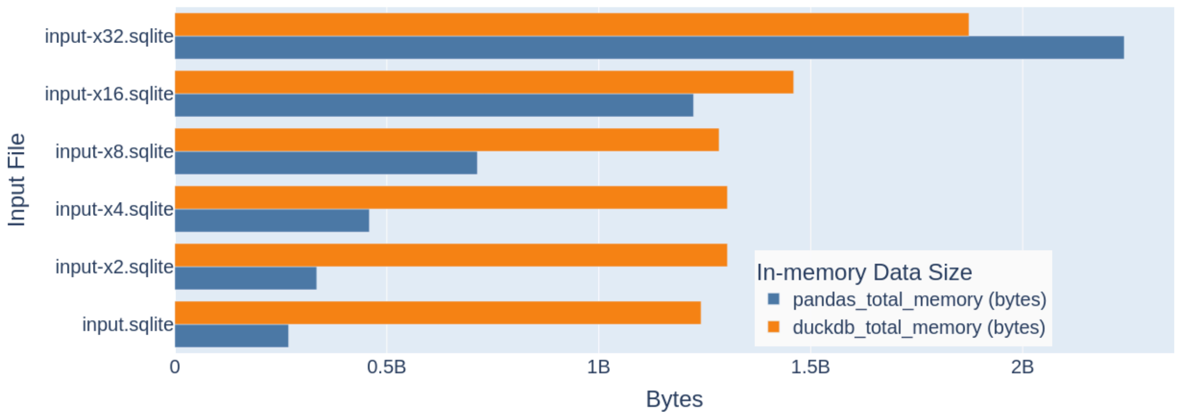


**Extended Note 2, Figure 4**. *Performance benchmarks of DuckDB and Pandas.* (Top panel) DuckDB required slightly more time than Pandas for join operations, but it surpassed Pandas in speed once input size exceeded ~30 MB (SQLite benchmark). (Bottom panel) DuckDB consistently used less memory than Pandas across larger datasets.

#### CytoTable MapReduce

CytoTable adopts a MapReduce paradigm to enable scalable and modular processing of large datasets. In the "map" phase, CytoTable processes data at the level of individual pages, allowing transformations, computations, or annotations to be applied independently and in parallel (**Extended Note 2, Figure 5**). This design not only optimizes memory usage but also lends itself naturally to distributed or batch processing frameworks. In the subsequent "reduce" phase, CytoTable recombines the processed pages into unified data structures. For example, pages may be concatenated to reconstruct the full table or joined with metadata and annotations to create enriched, analysis-ready datasets. This flexible approach enables CytoTable to handle large-scale data workflows efficiently.

| Original | Paginated | Changed | Reduced |
| --- | --- | --- | --- |
| "Data source" | Page or "Chunk" 1; Page or "Chunk" 2 | Modified Page or "Chunk" 1; Modified Page or "Chunk" 2 | "Reduced" data result (concatenated) |

**Original** – "Data source"

| Col_A | Col_B | Col_C |
| --- | --- | --- |
| 1 | a | 0.01 |
| 2 | b | 0.02 |

**Paginated** – Page or "Chunk" 1

| Col_A | Col_B | Col_C |
| --- | --- | --- |
| 1 | a | 0.01 |

Page or "Chunk" 2

| Col_A | Col_B | Col_C |
| --- | --- | --- |
| 2 | b | 0.02 |

**Changed** – Modified Page or "Chunk" 1

| Table_ID | Col_A | Col_B | Col_C |
| --- | --- | --- | --- |
| 123 | 1 | a | 0.01 |

Modified Page or "Chunk" 2

| Table_ID | Col_A | Col_B | Col_C |
| --- | --- | --- | --- |
| 123 | 2 | b | 0.02 |

**Reduced** – "Reduced" data result (concatenated)

| Table_ID | Col_A | Col_B | Col_C |
| --- | --- | --- | --- |
| 123 | 1 | a | 0.01 |
| 123 | 2 | b | 0.02 |

**Extended Note 2, Figure 5**. *MapReduce techniques enable mapped changes to take place on paginated sets which can be reduced to data summaries as concatenated tables.*

#### CytoTable has unified path handling for local or cloud-based data

CytoTable supports both local and remote data sources through a unified interface powered by the cloudpathlib library (**Extended Note 2, Figure 6**). This allows users to process data stored on Amazon S3, Google Cloud Storage, and Azure Blob Storage as if they were local files (leveraging commonly used paths denoted such as "s3://bucket/file"). For example, CellProfiler outputs hosted on public-facing S3 buckets can be read directly without authentication (for example, by leveraging "no-sign-request" configuration options).

For cloud-based SQLite sources, which require local access due to SQLite's file system constraints, CytoTable uses cloudpathlib's caching capabilities to download the database prior to processing. To avoid issues with system memory constraints in temporary directories, users may specify a custom cache directory (local_cache_dir). This balances the ability to use cloud-based paths even for data which may not inherently be streamable.

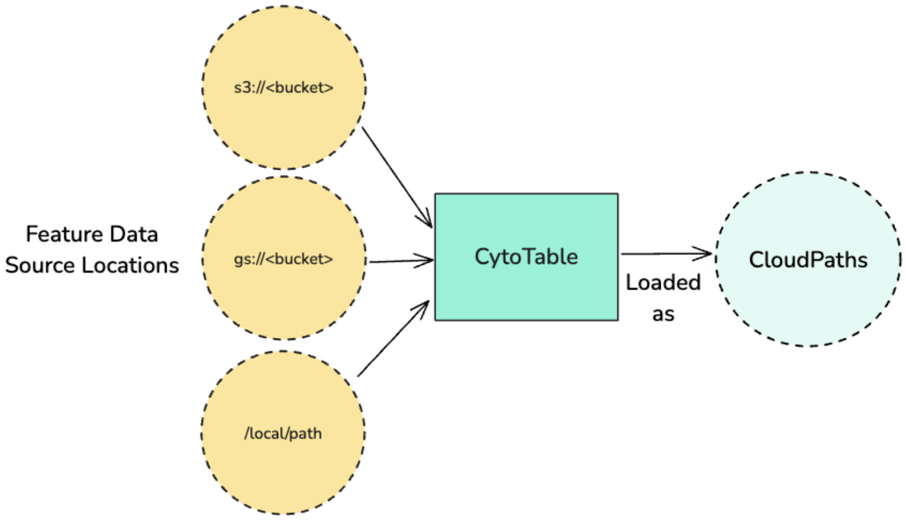


**Extended Note 2, Figure 6**. *Local paths and cloud-based paths can be handled as abstract CloudPaths within CytoTable making it easy for users to leverage datasets from many different locations without changing their code using recognizable filesystem patterns. Without CytoTable*

*and CloudPaths, scientific developers would need to implement more custom "stitch" code to stream or download cloud-based data so it may be treated as locally available data.*

### Extended Note 4 - CytoTable solves core image-based profiling challenges

#### SQLite type flexibility challenges

CytoTable uses SQLite as a source database format that is converted into more structured representations for downstream analysis. One of SQLite's distinguishing characteristics is its use of flexible typing, a system where column types are treated as type affinities rather than strict constraints. In this model, SQLite applies a best-effort mapping between the declared column type and the actual storage class of each value. In other words,  values in a single column can legally belong to different internal storage types.[53] For example, a column declared as REAL may contain both 0.01 (a floating-point number) and "value" (a string), corresponding to the REAL and TEXT storage classes, respectively. This flexibility can lead to challenges when consuming such data with systems that expect uniform data types.

Additionally, SQLite supports constraints like NOT NULL, which explicitly prevent the use of the NULL storage class, SQLite's marker for missing values. In practice, we encountered source tables where columns were constrained with NOT NULL but contained values such as "nan"—a string literal we inferred to represent numpy.nan[54], a NumPy standard in Python for null-like values in floating-point arrays. This kind of inconsistency is tolerated by SQLite, but creates friction when interoperability with strictly typed systems is required. In contrast, engines like DuckDB, as well as libraries like Apache Arrow and NumPy, enforce uniform types within columns or arrays. These systems do not permit mixed-type columns, making them incompatible with SQLite tables that include such heterogeneity.

To handle this mismatch, we implemented a workaround in CytoTable that dynamically constructs SQL queries using complex CASE expressions when exceptions related to this issue occur. These expressions detect and filter values based on SQLite's internal storage classes, allowing us to extract consistent, type-safe subsets of data. The NOT NULL constraint inhibited us from using potentially more efficient value replacement operations on the columns. While DuckDB performs most SQL operations in CytoTable are performed for its performance and analytical capabilities, these specific queries had to be executed within SQLite itself due to DuckDB's stricter typing enforcement. Although this solution was effective, it introduced significant engineering overhead and does not perform well due to the operating constraints. Based on this experience, we advise against using SQLite as a long-term data storage format for image-based profiling workflows, particularly because consistent data types are critical for performance, validation, and downstream tool compatibility.

#### Data type precision opportunities in image-based profiling

In image-based profiling workflows, numerical precision (particularly when storing floating point values) is sometimes an overlooked aspect of data integrity. Common formats such as CSV,

SQLite, and NumPy offer limited control over numeric precision and type representation. For example, NumPy arrays are typically constrained to fixed-precision types like float64, and SQLite's type affinity model treats numeric fields flexibly, often mixing strings and floats in the same column. These formats may silently truncate or round values, which can pose issues for reproducibility and downstream analysis, particularly when subtle differences in feature magnitudes are biologically meaningful.

In contrast, modern data systems such as DuckDB and Apache Arrow offer more explicit and robust support for high-precision decimals. As shown in the accompanying notebook (please see "explore_floating_point_precision.ipynb" notebook within CytoTable-benchmarks repository[35]), these systems allow for the definition and preservation of decimal types (e.g., `decimal128(17, 16)`), ensuring consistency across in-memory operations and persistent storage formats like Parquet. This level of precision could be important for HCI data where operations like normalization, aggregation, or phenotype scoring rely on subtle value differences. By preserving exact decimal representations, these platforms help maintain data fidelity and enable more accurate downstream modeling, making them increasingly attractive for use in open-source profiling workflows.

As tools in the profiling ecosystem evolve, attention to numeric precision could become a larger concern to help create opportunities for profiling. Choosing formats and engines that enforce or respect explicit numeric types (e.g., Arrow-backed DataFrames and DuckDB SQL engines) can standardize the representation of values across workflows and improve confidence in analytical results. CytoTable's support for Arrow and DuckDB aligns with this principle, ensuring that profiling outputs retain their intended precision across the full data lifecycle, from in-memory processing to persisted datasets.

#### Apache Arrow memory management considerations

Apache Arrow, through its Python bindings in PyArrow, employs a high-performance, off-heap memory architecture that enables fast, zero-copy data access across language boundaries. Instead of using Python's traditional object memory management system (i.e., the Python heap), PyArrow often allocates large contiguous blocks of memory through a native memory pool. These allocations happen outside the purview of Python's garbage collector (GC), which allows for efficient sharing of columnar data between Arrow, NumPy, DuckDB, and other systems without serialization overhead. The allocator (whether malloc, jemalloc, or mimalloc) can be controlled through the ARROW_DEFAULT_MEMORY_POOL environment variable, which influences how memory is reserved and reclaimed during processing.

While this memory model offers substantial performance benefits for large-scale data workflows, we observed complications when using PyArrow within multithreaded or multiprocessed environments during CytoTable development. Specifically, PyArrow-allocated memory appeared to not always be released in a timely manner, especially under high-throughput conditions involving large batches of Arrow Tables. In several instances, the Python GC struggled to detect and free unused memory buffers, likely due to the lack of tight integration between Python's GC and Arrow's native memory pools. This led to unexpectedly high heap memory usage within memory profile reports, and in extreme cases, appeared to lead to memory exhaustion. Apache Arrow's memory management differs from what is found within CPython and as a result likely is the cause for these discrepancies.

To mitigate these issues, we experimented with different memory allocators and batch sizes, and in some cases introduced manual deallocation patterns or forced GC cycles. We found that the choice of memory pool (controlled via ARROW_DEFAULT_MEMORY_POOL environment variable) could significantly impact both observed performance and memory stability. While PyArrow's design is well-suited for high-performance analytical computing, users integrating it into complex pipelines, such as those used in image-based profiling, should remain aware of these edge cases and monitor memory usage closely when scaling up to threaded or distributed workloads.
